# Empowering Integrative and Collaborative Exploration of Single-Cell and Spatial Multimodal Data with SGS

**DOI:** 10.1101/2024.07.19.604227

**Authors:** Tingting Xia, Jiahe Sun, Fang Lu, Yongjiang Luo, Yudi Mao, Ling Xu, Yi Wang

**Author notes:** These authors contributed equally to this work (Tingting Xia, Jiahe Sun).

## Abstract

Recent advancements in single-cell and spatial omics technologies have revolutionized our ability to capture multiple modalities data at a genome-wide scale within individual cells. However, visualizing these large-scale, high-dimensional, and complex datasets poses significant challenges. Here, we present SGS, a user-friendly, collaborative and versatile browser for visualizing single-cell and spatial multiomics data. SGS incorporates a novel genome browser framework, flexible visualization modules and a multi-panel adaptive communication mechanism to enable the synchronous visualization of diverse datasets. Notably, SGS empowers users with advanced capabilities for comparative visualization, through features like scCompare, scMultiView, and dual-chromosome module. Additionally, by adopting the “workstation concept”, SGS enables data fast visualization and collaborative exploration. We showcase the potential of SGS in the comparative visualization and coordinated exploration of mutlimodal data with two examples. SGS is publicly available at https://sgs.bioinfotoolkits.net/home.

## Main

Single-cell and spatial multimodal omics (scMulti-omics) technologies have empowered the measurement of diverse modalities^1,2^, including RNA expression, protein abundance, DNA methylation, chromatin accessibility, and spatial information at the cellular level^3–5^. The scMulti-omics has revolutionized our comprehension of gene regulation, epigenetic variations, protein expression dynamics, and cellular interactions, providing profound insights into complex biological processes^6,7^. Driven by innovative platforms, diverse tools and novel methodologies, single-cell and spatial omics research has experienced unprecedented growth in recent years, generating vast amounts of multi-omics data^8,9^. These multi-omics data exhibits substantial complexity^10^, characterized by the simultaneous presence of diverse and distinct feature sets within individual cells, such as (1) transcriptome and chromatin accessibility^11^; (2) transcriptome and DNA methylation^12^; (3) transcriptome and histone modifications^13,14^, etc. Moreover, multimodal data exhibits inherent abundant feature dimensions, spanning a wide range from hundreds of features for protein epitopes to hundreds of thousands for chromatin-accessible sites^15^. How to jointly visualize and collaboratively explore these multi-omics data of such high dimensionality, high noise and diversity poses significant challenges.

Several visualization tools have been developed to address these challenges^16,17^. Tools such as Cellxgene^18^, UCSC Cell Browser^19^, Loom-Viewer^20^, Loupe Browser^21^, ST Viewer^22^ and TissUUmaps3^23^, enable interactive exploration of single-cell features in lower dimensional UMAP or t-SNE spaces along with categorical, continuous and spatial dimensions, as well as feature annotation. However, these visualization tools were often designed for specific modalities or data types, such as scRNA or scATAC, which limit their capacity to handle the growing complexity of multimodal data and perform comparative visualization effectively. Researchers are often forced to utilize disconnected visualization tools to explore various aspects of multimodal data, which can be time-consuming. Several tools have been developed to bridge the gap in visualizing more complex multimodal data, including AtlasXplore^24^ and Vitessce^25^. However, it is important to acknowledge that these tools often require users with programming skills, making them less accessible to non-programmers. Furthermore, both tools have notable constraints in effectively visualizing epigenomic multimodal data. Overall, most existing visualization tools have limitations in installation convenience, multimodal integration, and comparative visualization highlight the urgent need for the development of new visualization tool (see **Supplementary Table 1** for a comparison with existing tools). It should be more accessible, versatile, and capable to facilitate in-depth interpretation of scMulti-omics data within a unified framework.

To address these limitations, we introduce the Single-Cell and Spatial Genomics System (SGS) as an innovative solution for enhancing the exploration of scMulti-omics data (**Figure 1 A**). SGS offers several key advantages compared to existing tools: (1) Multimodal integration and coordinate visualization. SGS offer two modules: SC (single-cell and spatial visualization module) and SG (single-cell and genomics visualization module), with adaptable interface layouts and advanced capabilities (**Figure 1 C**). Notably, the SG module incorporates a novel genome browser framework that significantly enhances the visualization of epigenomic modalities, including scATAC, scMethylC, sc-eQTL, and scHiC etc. To synchronous exploration of multimodal datasets with linked views across different panels, an adaptive communication mechanism was used bridges visualization component. (2) Comparative Visualization of different modalities and features: SGS offers dual-chromosome module, scMultiView, and scCompare functions to facilitate comparative visualization of cellular heterogeneity, gene expression, and genome-mapped signals of multimodal data. (3) Enhanced Accessibility and Collaboration. based on “data workstation” concept, SGS aims to empower research teams in swiftly establishing a visualization and sharing platform for collaborative data exploration without programming (**Figure 1 B**). (4) Compatible with diverse data formats and tools: SGS supports various data formats including AnnData/AnnData.zarr, MuData/MuData.zarr, and genome-mapped files (GFF, VCF, BED, HiC, Biginteract, Longrange, MethylC, Gwas, Bedgraph). The SgsAnnData R package enables seamless data format conversion with analysis tools like Seurat^26^, ArchR^27^, Signac^28^, and Giotto^29^ (**Supplementary Figure 8**).

**Figure 1:**
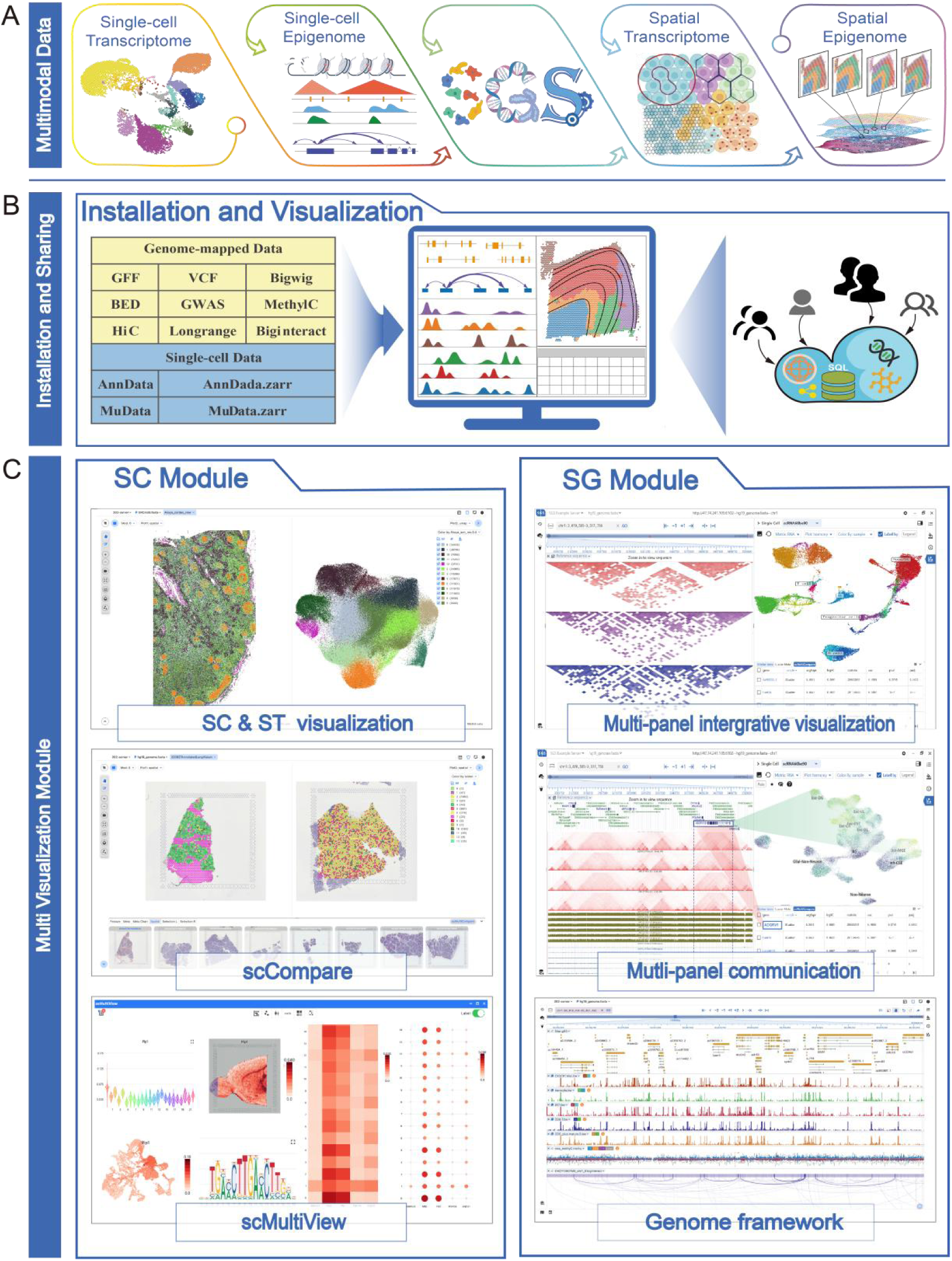
Overview of the SGS Browser. (**A**) SGS is a comprehensive visualization tool designed for single-cell and spatial multimodal data. (**B**) It supports various data formats, including AnnData/.zarr, MuData/.zarr, and genome-mapped files (GFF, VCF, BED, HiC, Biginteract, Longrange, MethylC, Gwas). The browser features an interface installation function and collaborative visualization of multi-omics data. (**C**) SGS offers two visualization modules: SC module and the SG module. The SC module provides a dynamic interface for exploring high-dimensional datasets, including core features such as 2D embedding plot, spatial multi-sample comparative visualization, and the scMultiView module for multi-gene comparisons. In contrast, the SG module introduces a novel genome browser that enhances the visualization of genome-mapped data. It integrates the genome framework with single-cell components and adaptive communication mechanisms to enable a coordinated multi-panel view.

In summary, the SGS provides a unified interface for visualizing scMulti-omics data, including genomics, transcriptomics, proteomics, and epigenomics data, at single-cell or spatial resolution. This comprehensive visualization browser facilitates the systematic interpretation of cellular heterogeneity, tissue organization, and biological processes.

## Results

### Graphical Installation and Collaborative Data Visualization

Efficient big data visualization is crucial for research teams. However, developing a new data visualization platform can be challenging and time-consuming for non-technical users. To address this, SGS adopts the “data workstation” concept and follows a “server-client separation” development strategy, enabling rapid develop of visualization platform for research teams.

First, SGS leverages Docker and Flutter technologies to achieve graphical one-click installation, avoiding the complex software configuration and web service deployment. This makes SGS compatible with multiple systems, including Linux, Windows, MacOS, and Android. With just a single click, SGS automatically pulls the necessary images and runs a Docker container with all required software. Users can initiate the installation process by providing essential information such as server IP and password. The progress can be monitored in real-time through a progress bar (**Figure 2 A, B**) (**Supplementary Figure 1**). Once the installation is complete, users can batch upload and visualize various data files, including GFF, VCF, Bigwig, AnnData and MuData (**Figure 2C**).

**Figure 2:**
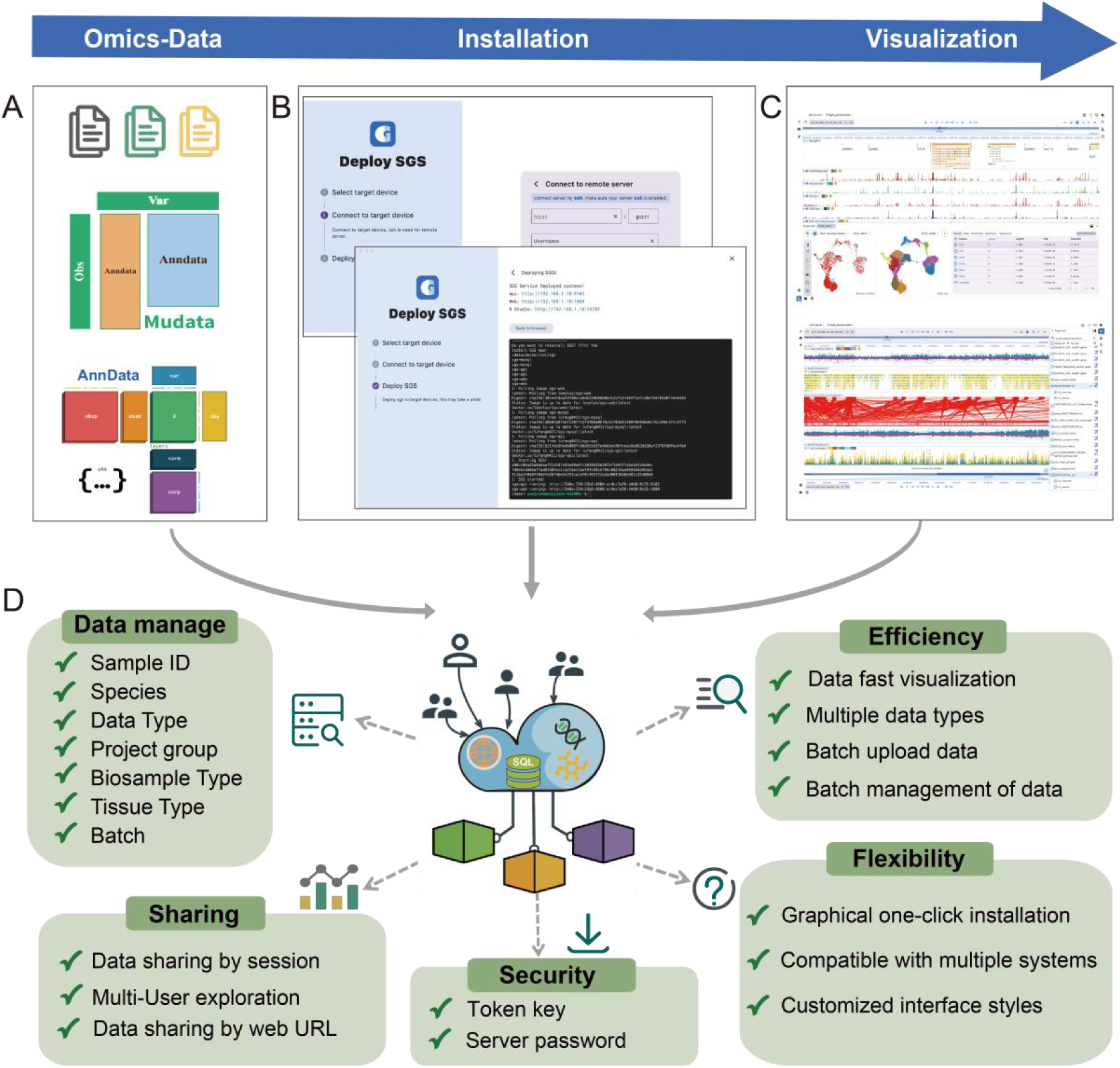
Graphical Installation and Data Visualization. (**A**) SGS supports a wide range of omics data formats, including AnnData, AnnData.zarr, MuData, MuData.zarr, GFF, VCF, Bigwig, Bed, Gwas, MethylC, Hic, Longrange, BigInteract, etc. (**B**) SGS offers a user-friendly graphical installation interface, guiding users through the installation process with progress bars and prompt boxes. (**C**) SGS provides extensive visualization capabilities and offers diverse interface layouts. (**D**) SGS incorporates collaborative visualization features that optimize data batch management, data sharing, security and privacy, visualization efficiency, and software flexibility

Beyond its rapid setup capabilities, SGS provides a comprehensive suite of features designed to enhance collaborative data exploration and visualization sharing: (1) Real-Time Collaboration: SGS offers a unified client interface that can connect to different server instances, eliminating the complexities of large data migrations. Multiple users can work on the same dataset, with real-time updates and synchronization. This fosters a collaborative environment where insights can be shared and discussed instantly. (2) Seamless Data Sharing: SGS allows users to share view sessions or web URLs. This feature makes it easy for team members to access and collaborate on data visualizations. (3) Advanced Data Access Controls: Recognizing the importance of data security, SGS implements data access control mechanisms. It utilizes token-based authentication to restrict data access to authorized users or teams, ensuring the security of sensitive information (**Figure 2 D**).

In summary, SGS has brought significant enhancement to research teams in terms of simplifying deployment, improving data uploading and management efficiency, and protecting data security. Making it an valuable tool for scientific research teams aiming to efficiently visualize and share large datasets..

### Single-Cell and Spatial Omics Data Visualization

#### SC Module

The SC module is specifically designed for comparative exploration of non-genome-mapped scMulti-omics data, such as scRNA, ST and scProteomics etc. The SC module supports embedding plot (from UMAP, t-SNE or PCA) and visualization of tissue slices (**Figure 3 A (a)**). To allow users to more intuitively view cell annotation information, the SC panel supports overlays various metadata onto the scatter plot, including cell type annotation, clustering information, and sample details (tissue, donor, and other clinical features) (**Figure 3 A (b)**). Users can use the slider to flexibly adjust the size of the point, transparency, and brightness of the H&E image. The SC component also supports zooming, dragging, selection and other operations to explore the region of interest (RIO). Additionally, users can conveniently click or search gene to query differences in gene expression across cell types. At the bottom of the main interface prominently displays essential information, including tissue slices, meta chart, marker gene table, metadata and cell subset (**Figure 3 A (c-e)**). These features enable examining cell annotations, exploring marker expression differences, comparing cell compositions across tissues, and visualizing multiple spatial slices.

**Figure 3:**
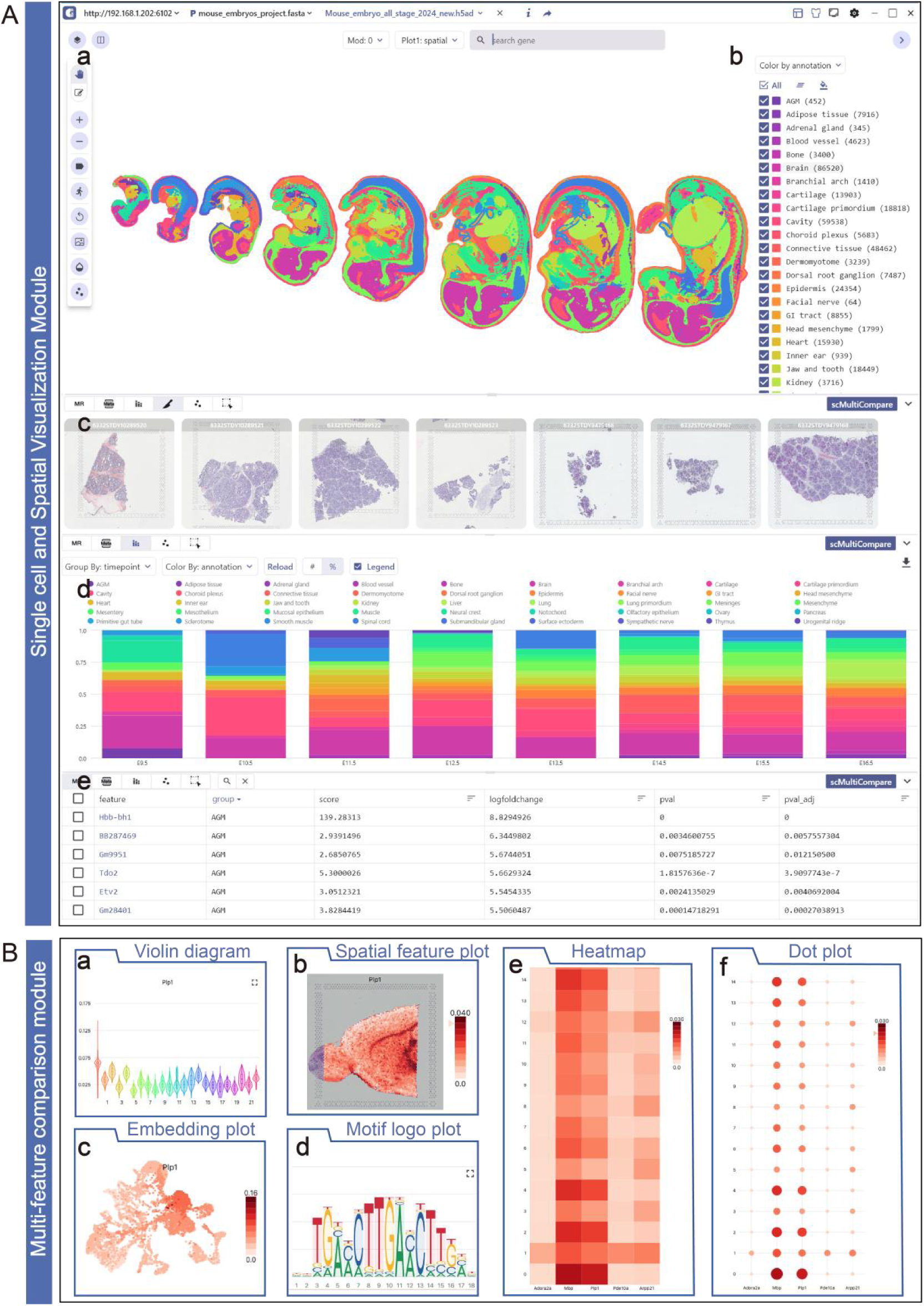
Key Visualization Features of the SC Module. (**A**) The main panel of the SC module supports the display of embedding and annotation information of mouse embryo development (**a, b**). The Spatial Slice tab presents all tissue slices from human lung ST multi-sample datasets (**c**). The Meta Chart tab showcases cell proportions using stacked bar charts (**d**). The marker feature table provides marker information, including gene name, group, P-value, log fold change, and other relevant details (**e**). (**B**) The scMultiView module offers a diverse range of visualization charts. These include violin plot (**a**), spatial feature plot (**b**), embedding plot (**c**), motif logo plot (**d**), heatmap (**e**), dot plot (**f**).

In various research studies, the comparative function is crucial. Researchers can compare gene expression patterns across multiple sample groups (e.g., case and control groups, male and female groups, young and middle-aged and elderly groups). The scCompare function enables users to easily compare specific cell type annotations and gene expressions through a simple click of the “Split View” button. Additionally, it allows side-by-side views for comparing gene expression patterns or tissue slice characteristics across multi sample in ST experiments.

Furthermore, to further enhance the comparative visualization capability of multiple genes, we have developed the scMultiview component. It offers a wide range of visualization options, including violin plot, spatial feature plot, embedding plot, motif logo plot, heatmap, and dot plot (**Figure 3B (a-f)**). Among these, annotation-based embedding plot can identify differentially expressed genes within specific cell types. The spatial feature plot illustrates the spatial expression characteristics of genes. Gene expression violin and box plots displayed the expression differences between cell types, facilitating comparisons across groups. motif logo plot reveal shared motifs of transcription factor binding sites, aiding in the elucidation of gene regulatory networks. When multiple genes are selected, users can generate heatmap or dot plot to visually display the expression differences, assisting in cell type annotation.

In general, SC visualization module provides a dynamic interface for exploring high-dimensional datasets. Its core features include single-cell embedding plot, metadata visualization, comparative visualization of multiple spatial samples or genes. These functionalities assist in identification of differentially expressed genes, cell type annotation, exploration of cellular heterogeneity, and understanding complex biological processes at the single-cell level.

The complexity of single-cell and spatial epigenomic omics data involving multi-layers information, including 3D genomic architecture, histone modifications, and chromatin accessibility^30^. To achieve a comprehensive understanding and extract valuable insights from these datasets, need the integration of information from both genomic and single-cell resolution levels. Here we have developed the SG Visualization Module, which integrates the visualization capabilities of genome browser framework and single-cell component (**Figure 4 A (a,d))**. To improve the compatibility of genome-mapped data and enhance the scalability, customizability, and interactivity of the SG visualization module, we have developed a novel, flexible, and scalable genome browser framework. This framework enables the visualization of various genome-mapped signals, including gene annotations, epigenome modification signals, genetic variations, and eQTLs, among others. It ensures broad compatibility by supporting popular genome-mapped file formats, including GFF, VCF, Bigwig, BED, MethylC, HiC, Biginteract, and Longrange. Additionally, it not only supports commonly used operations in existing genomic data visualization tools but also provides many novel features to facilitate integrative exploration of genome-mapped signals, including optimized feature display module, dual chromosome visualization module, multi-track group, etc (**Supplementary Figure 4**). The SGS dual-chromosome visualization module provides a powerful visual comparison of single-cell and spatially epigenomic multimodal signals. By utilizing a double chromosome display strategy, this module presents several advantages. Firstly, the top and bottom coordinates can cover different genomic regions, enabling the visualization of single-cell long-range interactions (**Figure 4B (a,b,c)**). Secondly, users have the flexibility to independently shift or zoom these coordinates region. This capability facilitates the comparative visualization of cell type-specific epigenomic signal differences in multiple regions simultaneously, enhancing our understanding of cell-specific regulatory patterns. Users can freely arrange and customize these components according to their needs, enabling synchronized views of multimodal datasets or different views of the same data modality (**Supplementary Figure 5 (A,B))**. The integrative panel design of the SG Visualization Module holds potential for uncovering intricate relationships and interactions among diverse types of molecular information within distinct omics layers.

**Figure 4:**
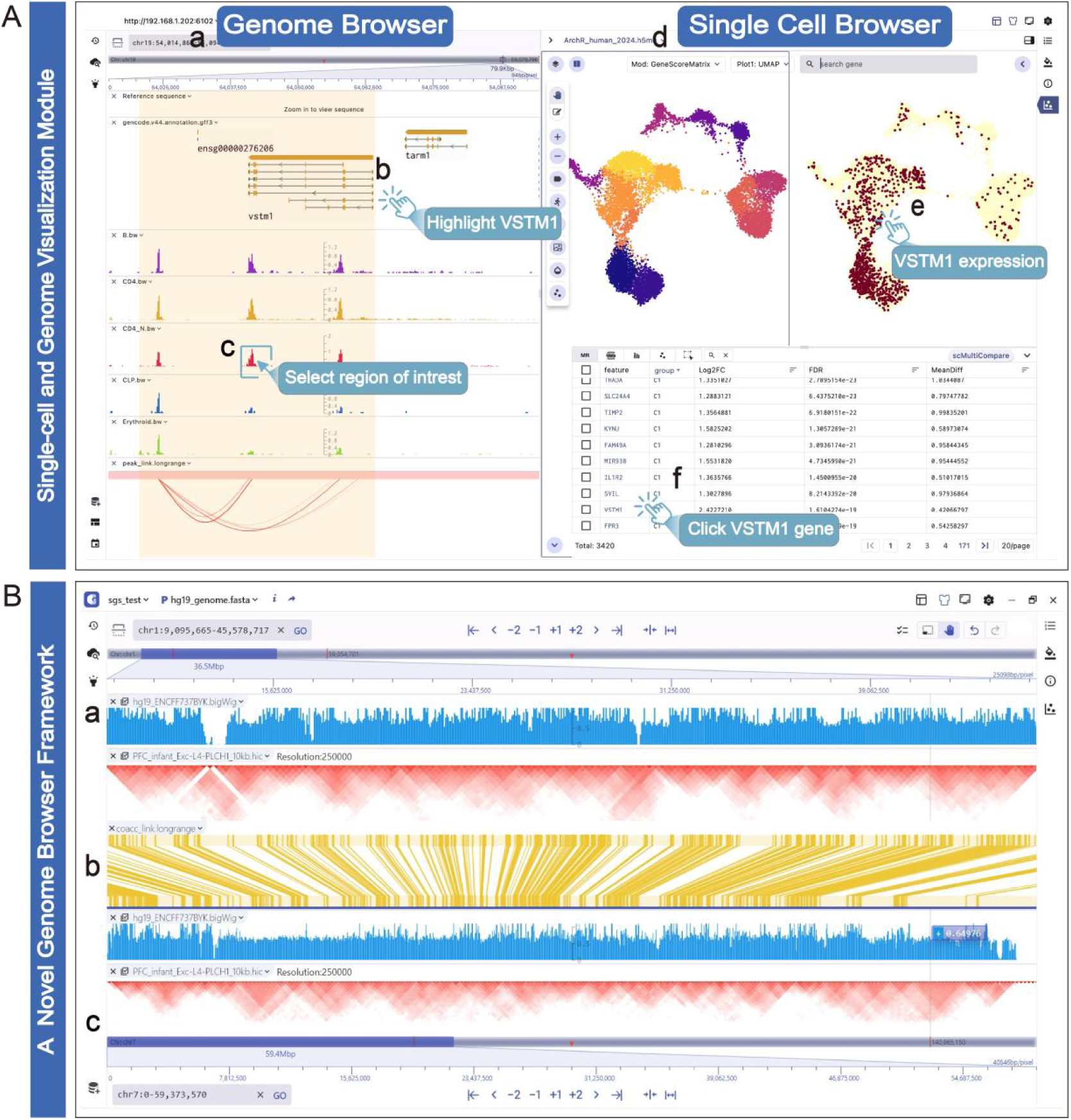
Main Features of SG Module. (A) The SG visualization module displays human hematopoietic scATAC datasets with two main components: the genome framework (left) and single-cell components (right) (**a,d**). The genome browser shows genomic tracks for the *VSTM1* gene, including gene structure and chromatin accessibility signals for PreB, Mono, CD4 N cells, and other types. It also displays co-accessibility links between peaks and genes. Users can navigate, highlight, zoom, and select specific regions and features (**b,c**). The single-cell panel features a split-screen with cell embedding plot and *VSTM1* gene expression distribution (**e**). Clicking on marker genes in the table allows exploration of gene activity scores and genome-mapped signals (**f**). **(B)** The SGS double-chromosome visualization mode enables easy comparison of genome-mapped data across different chromosome. The top (**a**) and bottom (**c**) panels display human Bigwig track and Hic interaction track on chr1 and chr7, respectively. The Bigingteract track in the center panel shows interaction derived links between the genomic regions (top coordinate) and other genomic regions (bottom coordinate) (**b**). The strengths of the genomic interactions are plotted in color scale, with red being strongest. Users can freely switch between the top and bottom visualization areas to explore the data.

To gain a deeper understanding of the intermolecular dynamics between epigenome modifications and cellular heterogeneity, we have defined an adaptive communication mechanism further enhance the integration of these two components. Currently, the mechanism is primarily based on genomic feature ID like peaks, genes and sc-eQTL as the communication protocols to achieve the coordinate view between genome browser framework and single-cell panel **(Supplementary Figure 6)**. When users click on these features in any interface, the visualization panels automatically query, navigate, and render the dynamic changes of these features at both the genome and single-cell levels. For example, The SG panel in the figure displays genomic signaling and cell embedding plot of human hematopoietic datasets from previous studies^31^. when a user selects an interesting feature (such as the *VSTM1* gene) in the genome browser framework, an event listener in the single-cell panel receives the gene feature ID and synchronously highlights its expression distribution in different cell types. By clicking on the *VSTM1* gene in the single-cell panel’s marker table **(Figure 4A (f))**, users can immediately observe their expression patterns across different cell types **(Figure 4A (e))** and navigate to the gene region to explore the distribution of epigenomic modification signals **(Figure 4A (b,c))**, such as chromatin accessibility and peak-to-gene links, in the genome browser framework. This mechanism facilitates synchronous exploration of identical data types with linked views, allowing multimodal datasets to be displayed across different browser panels.

Overall, the SG visualization module offers a scalable solution for integrating an ever-expanding volume of epigenomic multiomic datasets. By integrating genome browser framework and single-cell panel, SG enables synchronous exploration and comparisons of various datasets (e.g., snATAC, scM&T-seq, snm3C-seq, and sce-QTL) across linked panels through multi-panel adaptive communication mechanisms.

### User Case

To showcase the advantages of using the SGS for integrative single-cell and spatial multi-omics data exploration, we present two distinct examples of data integration.

As the first example, we show how SGS supports the synchronized visualization of Single-nucleus Methyl-3C Sequencing (sn-m3C-seq) data encompassing DNA methylation and 3D chromatin conformation during the development of the human frontal cortex (PFC) and hippocampus (HPC)^32^. Figure 5A displays two different views that integrate features selected in the SG visualization module: (1) The top view of genome browser displays the variation track of *rs500102*, positioned near the promoter region of the *RORB* gene (**Figure 5A (a)**). Below this view, the panel showcase the CG methylation signals alongside Hic interaction track of specific cell types (e.g., PFC 2T RG-1; PFC adult L1-3 NRXN2; PFC adult L4-5 FOXP2; PFC 2T eMGE; PFC adult MGE-ERBB4; PFC adult ODC) (**Figure 5A (b)**). (2) The single-cell panel showcases the cell atlas of sn-m3C-seq data obtained from 13 developmental adult PFC and 9 HPC samples (**Figure 5A (c)**), which provides an overview of the distribution of 10 primary cell types and CG methylation patterns of *RORB* gene (**Figure 5A (d)**). Users can select a key feature in the single-cell marker table to directly navigate to the corresponding region in the genome browser to obtain multi-omics layer information through the adaptive communication design. Of particular interest, by clicking on the *RORB* gene within the marker gene table, users can navigate to the corresponding region in the genome browser. This allows for the observation of the noticeable enhanced chromatin interaction strength specifically in the adult PFC L4-5 FOXP2 cell population within the *RORB* region, accompanied by a decreased CG methylation signal compared to other cell populations. Alternatively, by selecting the *RORB* gene in the genome browser, users can investigate the heterogeneity in CG methylation signal distribution among various cell populations. Notably, a significant decrease is observed especially in excitatory neurons within the PFC L4-5 FOXP2 cell cluster, which is consistent with previous research findings.

**Figure 5.**
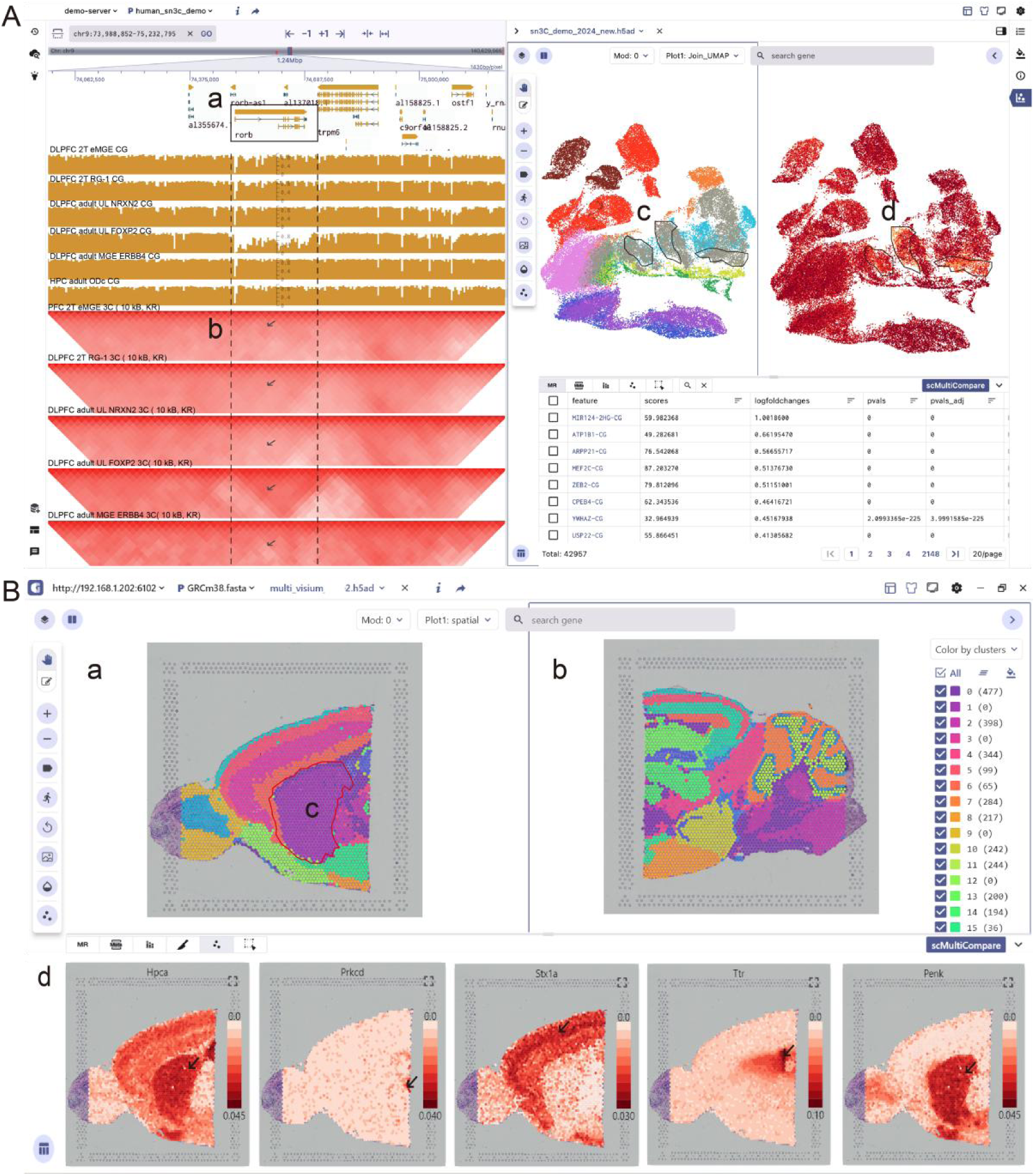
User case. (**A**) The main panel illustrates the single-cell epigenomic multi-omics data (sn-m3C-seq) capturing DNA methylation and 3D chromatin conformation during human frontal cortex (PFC) development. The left genome browser displays genome annotations in the region near the *RORB* gene, along with HiC and CG methylation signals for specific cell types: 2TRG-1, adult L1-3 NRXN2, adult L4-5 FOXP2, 2T eMGE, adult MGE-ERBB4, and adult ODC (**a,b**). The single-cell panel on the right presents the cellular profile of the PFC and a scatter plot showing the expression of the *RORB* gene (**c,d**). (**B**) The SC module mainly shows the 10x visium data of the anterior and posterior brains of mouse. The split screen of the main panel shows the spatial distribution difference of 14 cell clusters in the anterior and posterior brain tissues (**a,b**), and cluster1 is mainly exist in the posterior brain tissue (**c**). The spatial feature plots of marker genes including *STX1A* (cortext marker), *Prkcd* (thalamic marker), *HPCA* (hippocampal marker), *Ttr* (choroid plexus marker) are displayed at the bottom of the panel (**d**).

The second example showcases the comparative visualization of 10x Visium multi-sample data from the anterior and posterior regions of the mouse brain using SGS^33^. We utilized the scCompare function of the SC panel to present the spatial context of distinct tissue slices in side by side ((**Figure 5B (a,b)**). As illustrated in **Figure 5B (c)**, cell cluster1 specifically exists in the posterior brain tissue section. Furthermore, users can also utilize this function to observe the expression patterns of target genes in different sample slice regions. For example, our observations reveal that genes such as *STX1A, Prkcd, HPCA,* and *Ttr* exhibit specific expression in the cortex, thalamus, hippocampus, and choroid plexus regions, highlighting the spatial heterogeneity of gene expression within tissues. The bottom panel of the SC module shows thumbnails of multiple sample slices. Users can select any sample slice from the bottom panel and quickly identify the differences in cell proportions by clicking the “metachart” button to generate a stacked bar chart. For example, using sample ID as the grouping feature, we showcase the proportion differences of cell clusters across anterior and posterior brain tissues (**Figure 5B (d)**). Compared to the posterior brain sample, we observe that the anterior brain sample has a lower proportion of cells in cluster 0, while a higher proportion of cells in cluster 2 **(Supplementary Figure 2)**. For batch comparison visualization of multiple genes, users can select gene sets in the marker table or through the search box, and quickly compare and visualize them using the scMultiView component. For example, we performed batch comparison visualization for genes (*Hpca, Prkcd, Stx1a, Ttr, Penk*) with region-specific expression. Their expression features are displayed using various visualization charts such as scatter plot, violin plot, dot plot, or heatmap **(Supplementary Figure 3)**. This facilitates cell type annotation and in-depth exploration of heterogeneity in gene expression.

## Discussion

Comprehensive visualization of single-cell and spatial multi-omics data is crucial for investigating the intermolecular dynamic between epigenomic gene regulation and transcriptomic/proteomic gene expression within individual cells across development, ageing and disease. This have led to the generation of large-scale, high-dimensional, and complex multimodal data, posing challenges for scMulti-omics data visualization. In this paper, we introduce SGS, which provides a unified interface for the joint visualization of multi-omics data, including genomics, transcriptomics, proteomics, as well as single-cell or spatial resolution epigenomics. It offers modular panels for heterogeneous data types, combined with multiscale interactive navigation and collaborative exploration. SGS have many advantages and novel features: Firstly, Equipped with its graphical installation and “data workstation” concept, SGS effectively reduces the learning curve, enabling researchers to quickly install and engage in collaborative data visualization tasks. Secondly, SGS supports interactive visualization of single-cell clustering and spatial tissue slice. By leveraging the scCompare function and scMultiView module, SGS offers rich plotting capabilities, fostering comprehensive comparisons across single-cell and spatial samples, as well as multiple genes. Furthermore, to meet the demands of visualizing complex epigenomic multimodal data, we have developed a novel genome framework. This framework accommodates various data formats, provides multiple visualization features, and incorporates dual-chromosome visualization capabilities to enhance the comparative visualization of genome-mapped data. By further integrating single-cell component and genome browser framework within the unified panel, SGS enables to integrated visualization of diverse data types. The synchronized response design between these panels holds significant value by enabling integrated visualization, cross-modal insights, interactive exploration, and enhanced data interpretation.

We demonstrate SGS’s ability to visualize multimodal data through two case studies. The first case study highlights the synchronized visualization and cross-modal integration of sn-m3C-seq acquired from the PFC and HPC. By integrating gene annotation, SNP variation, CG methylation, 3D chromatin interaction, and single-cell views, SGS provides a comprehensive perspective for in-depth visual exploration of epigenomic multimodal data, facilitating research to investigate the diseases like schizophrenia. In the second case study, we utilized SGS to perform comparative visualization of complex multi-sample spatial data obtained from mouse brain. SGS facilitated intra-group and inter-group visual comparisons through features such as scCompare, scMultiView, and MetaChart. These functionalities provide valuable insights into the colocalization of cell types and cell states, spatial covariance of gene expression changes, and defining cellular tissue niches and how they change in disease.

With the upcoming massive wave of new and diverse scMulti-omics datasets, on the foundation of visualizing different modalities within the unified interface, the deeper integration of data features from multi modalities often requires advanced integration algorithms. SGS currently offers an appealing solution through its multi-panel integration and adaptive communications. To further enhance the exploration of high-dimensional data modalities, our future focus lies in the combination of advanced multimodal integration algorithms such as scAI^34^, MOFA+^35^, totalVI^36^, MAESTRO^37^, STvEA^38^ and others into SGS. Besides, as spatial sequencing technologies continue to advance, the visualization of data needs the extraction of signals from numerous image layers, each representing different fluorescent channels collected during various staining rounds. To enhance the ability to view individual molecular labels and effectively handle multi-layered image data, our efforts will be directed towards optimizing the rendering of imaging-based spatial transcriptomics^39^ (IST) data using the OME-NGFF (Open Microscopy Environment-Next-Generation File Formats) file format^40^.

In summary, the Single-Cell and Spatial Genomics System (SGS) empowers researchers with a powerful omics browser, unlocking novel insights from scMulti-omics data. With its exceptional flexibility, scalability, and accessibility, SGS plays a crucial role in advancing our understanding of differentiation trajectories, the underlying gene regulatory networks, cell-to-cell interactions, microenvironmental spatial organization, cellular lineages, and clonal dynamics. The platform holds promise for future advancements, accommodating new data modalities and integrating diverse omics datasets, further propelling scientific discoveries in the field.

## Methods

### Software Architecture and Implementation

The SGS browser adopts a front-end and back-end separation architecture. The software is divided into distinct layers, including the data layer, service layer, API layer, and client layer. The Flask-based service layer handles data parsing and processing. It efficiently handles data-related tasks within the SGS software. On the other hand, the client layer utilizes the Flutter framework to emphasize interactive visualization. The API layer mainly used for the communication between the service layer and the client layer. This enables seamless data exchange and integration between the different layers of the SGS browser (**Figure 6 B**).

**Figure 6:**
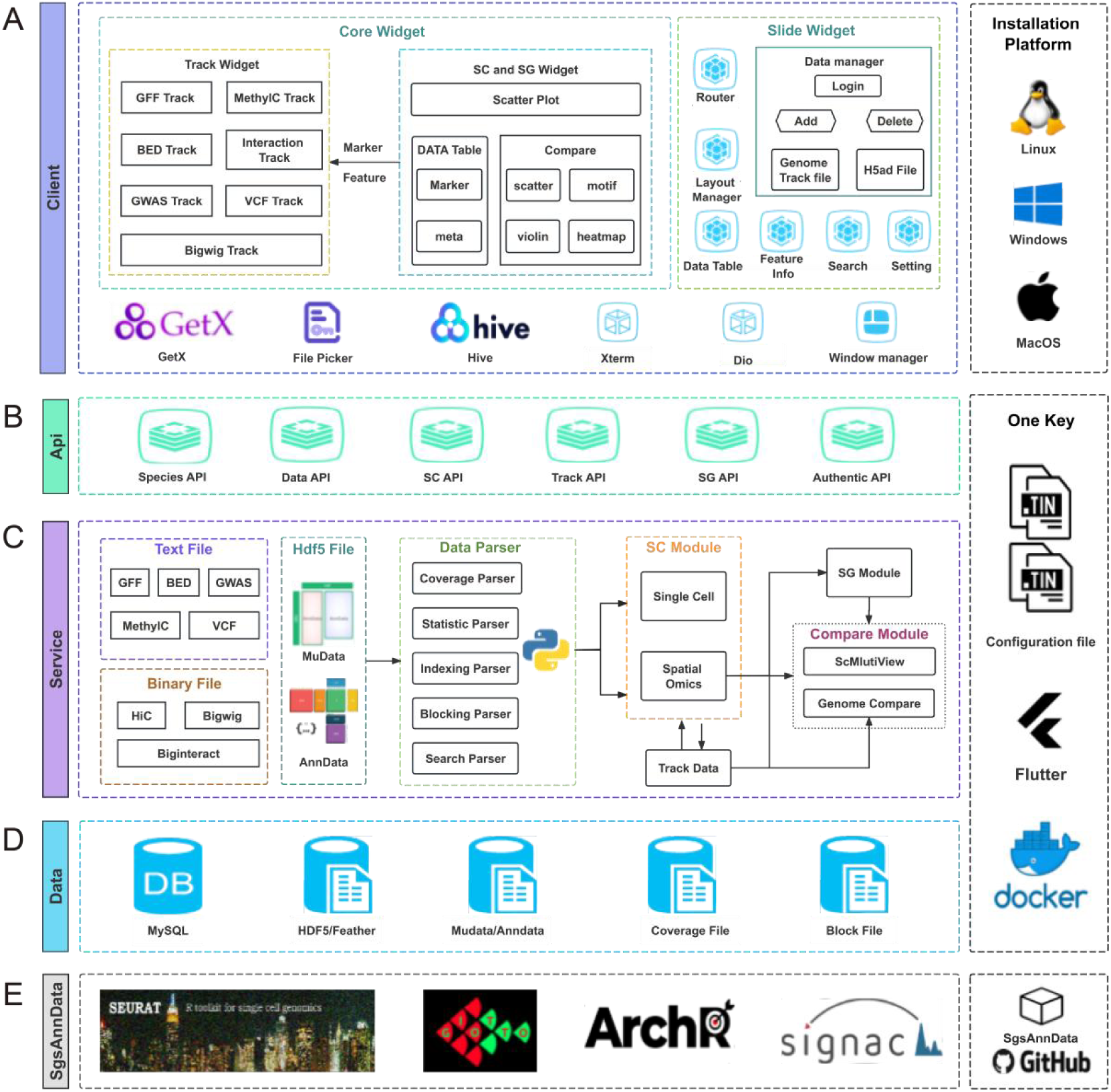
The framework of SGS. (**A**) The client development utilizes the Flutter framework, primarily consisting of Core Widgets and Slide Widgets. The Core Widgets include Track Widgets and SC/SG Widgets. This client is compatible with multiple platforms, including Linux, Windows, and MacOS. (**B**) The API layer defines a series of front-end and back-end communication interfaces, including Species API, Data API, SC API, Track API, SG API, Authentic API. (**C**) The server-side development primarily employs the Flask framework, encompassing the Data Parser, Single-cell and Spatial Modules (SC Module), Single-cell and Spatial Genomics (SG Module), and Comparison Module. (**D**) The data layer defines storage strategies for different types of data. SGS utilizes Docker technology to encapsulate back-end business logic and achieves GUI-based installation in conjunction with the Flutter framework. (**E**) SgsAnnData is an R package used for file analysis and AnnData data format conversion in tools such as Seurat, Giotto, ArchR, and Signac.

### Client Design

The client of the SGS browser utilizes the Flutter (version 3.16) framework, known for its robust cross-platform capabilities, enabling it to run seamlessly on various platforms such as macOS, Windows, and Linux. Leveraging Dart as the primary programming language, we utilize its rich functionality and extension library support to implement the features and interface of the SGS browser client. The SGS client mainly comprises two major Widgets: Core Widgets and Side Widgets (**Figure 6 A**). These components cater to different interaction requirements and contribute to the overall user experience. Core Widgets consist of three parts: genome browser widget, SG widget, and SC widget. On the other hand, Side Widgets primarily include Data Table Widgets, Search Widgets, and Feature Info Widgets, providing auxiliary functions for region highlighting, track list operations, theme, and layout settings.

### Server Design

The back-end using Flask (version 3.02) framework, is famous for its lightweight, provides system with a higher response speed and lower memory consumption and maintenance costs. The back-end server is primarily divided into four main components: file type identification and management, data parsing and calculation, data relationship mapping, and module communication.

Moreover, it utilizes a multi-threaded approach to facilitate concurrent data processing, enabling batch data uploads and enhancing data addition efficiency (**Figure 6 C**). For efficient data management and storage, we have employed MySQL (version 8.0) as our preferred solution. SGS leverages advanced techniques such as memory mapping and index building to optimize the performance of rapid queries, particularly in track transformation and the scaling of extensive datasets (**Figure 6 D**).

### Single-cell and Spatial Multi-omics Visualization Component

#### A Novel Genome Browser Framework

The genome browser framework supporting popular data formats such as Bigwig, Biginteract, GFF, VCF, MethylC, BED, Longrange, Gwas, and Hic. We implemented specific parsing strategies for binary and text file formats. For text formats (e.g., BED, VCF, GFF), we utilized tabix (version 1.7-2) to retrieve region-specific data. Binary formats (e.g., Bigwig and Hic) were parsed using pybigwig (version 0.3.22) and straw (version 0.1.0). Data storage employed the feather memory mapping technique for improved efficiency.

We simplified front-end visualization by implementing a universal Track Widget parent class. Customized SGS chart libraries support various track display styles including bar charts, gene structures, area charts, and heatmap. Different visualization levels were defined for feature presentation at varying zoom levels. Density calculation was performed at the chromosome scale, while data blocking was employed at the second and third zoom levels to enable efficient rendering and switching. Techniques such as preloading view data, dynamic retrieval, and memory management minimized storage consumption and ensured rapid feature rendering and zooming at different visualization levels.

#### Single-cell Component

The single-cell component includes the SC and SG widgets for visualizing cell embeddings and spatial tissue slice. We use AnnData (version 0.10.6) and MuData (version 0.2.3) for data storage, and the zarr format for large-scale multimodal data, which supports chunking and compression. Furthermore, to optimize marker feature queries, we utilize the N-gram language model [https://github.com/joshualoehr/ngram-language-model] which is a language model that utilizes n-grams to capture the contextual dependencies between words in a given datasets. To optimize rendering speed and minimize memory consumption, we employ techniques such as cell grouping rendering, gradient interval division, and random sampling. For spatial transcriptomic data, precise alignment of spatial coordinates with tissue slices is ensured by using scaling factors for different image resolutions. Additionally, a caching mechanism preloads spatial slice information, enabling swift loading and efficient visualization for comparing multiple samples. The scMultiView Widget supports various visualization plots, including violin plot, box plots, heatmap, and dot plot, providing comprehensive insights into single-cell transcriptomic data

#### Multi-Panel Adaptive Communication

We developed a multi-panel communication mechanism that integrates and coordinates single-cell, spatial, and genome visualization panels, enabling data exchange, collaborative exploration, rapid navigation, and highlighting across variant components. The system consists of a unified communication protocol, core communication engine, and functional module components. Each functional module component includes communication triggers and listeners, while the unified communication protocol defines data transmission methods and network connection rules.

The core communication engine coordinating communication signals between the genome browser framework and single-cell component. It manages data transmission, error handling, connection management, and security authentication to ensure effective links between different data modules. The Communication triggers enable user interactions, such as click in the single-cell panel to select marker features (such as genes, peaks, and sc-eQTLs). These actions are then transmitted to the core engine, which subsequently directs them to the communication listener within the genome browser framework. Upon receiving the signal, the listener swiftly retrieves and locates the position of the marker feature, synchronously highlighting the region and displaying other genome modification signals such as methylation signals, Hic chromatin interaction intensities, etc.

### SgsAnnData

Mainstream software tools such as Seurat, Scanpy, ArchR, Signac, Giotto, and Squidpy offer extensive capabilities for in-depth analysis of single-cell and spatial multi-omics data. To compatible with the output of these tools, we have developed the SgsAnnData R package (**Figure 6 E**). This package converts the data structures of these software into the widely compatible AnnData formats. Our comprehensive SgsAnnData tutorial [https://github.com/bio-xtt/SgsAnnDataV2/tree/main] provides detailed guidance on using SgsAnnData to convert outputs from different software into input data formats of SGS (**Supplementary Figure 9**).

### Demo Data Preparation

#### Mouse Embryo Spatial Transcriptomics Data

The mouse embryo developmental spatial transcriptomics example data used in this study were obtained from the research of Chen A, Liao S et al^41^. The data were primarily downloaded from the website: https://db.cngb.org/stomics/mosta/.

#### Human Lung Spatial Multi-slice Data

The human lung spatial multi-slice example data used in this study were sourced from the research of He P, Lim K et al^42^. The data were primarily downloaded from the website https://genome.ucsc.edu/s/brianpenghe/scATAC_fetal_lung20211206.

#### Human Hematopoietic Stem Cell scATAC Data

Initially, we downloaded the human hematopoietic stem cell fragment files^31^. Subsequently, we use ArchR and follow the tutorial guidelines [https://www.archrproject.com/bookdown/index.html#section] with default parameters. we performed analyses such as doublet removal, single-cell clustering and cell type identification, unified peak set generation, cellular trajectory identification, DNA element-to-gene linkage, and transcription factor footprinting. The file format conversion was accomplished using the ArchrToAnndata function of the SgsAnnData (version 0.3.0) package with default parameters.

#### Human PFC and HPC sn3C-seq Data

The human PFC and HPC sn3C-seq example data used in this study were obtained from the research of Matthew G. Heffel et al^32^. The data were downloaded from the website: https://cells.ucsc.edu/?ds=brain-epigenome+human-brain-m3c.

#### Mouse Anterior and Posterior Brain Spatial Multi-sample Data

We downloaded the mouse anterior and posterior brain data from the 10x Genomics website [https://www.10xgenomics.com/datasets?menu%5Bproducts.name%5D=Spatial%20Gene%20Expression] and performed data quality control, spatial integration of multiple samples, dimensionality reduction, clustering, and cell type annotation using Seurat (version 4.3), following the official tutorial. Finally, we transformed the data into adata using the SeuratToAnndata function of SgsAnnData (version 0.3.0).

#### Human oneK1K sc-eQTL Data

The human oneK1K sc-eQTL data used in this study was sourced from Seyhan Yazar et al^43^. The single-cell data was primarily obtained from the website: https://cellxgene.cziscience.com/collections/dde06e0f-ab3b-46be-96a2-a8082383c4a1, while the eqtl gwas data was downloaded from https://onek1k.s3.ap-southeast-2.amazonaws.com/onek1k_eqtl_dataset.zip.

#### ME11 Spatial-ATAC-seq Data

The mouse embryo spatial-ATAC-seq (1 1.5 days) data from Deng Y, Bartosovic M et al^44^. study was downloaded from https://www.ncbi.nlm.nih.gov/geo/query/acc.cgi?acc=GSE171943. We reproduced the relevant analysis results following the analytical pipeline provided by the authors [https://github.com/dyxmvp/Spatial_ATAC-seq]. The file formats were converted using the ArchrToAnndata function of SgsAnnData (version 0.3.0) and mudata (version 0.2.3) packages.

### Availability of Code, Software and Documentation

The code of SGS is available at: https://github.com/fanglu0411/sgs?tab=readme-ov-file. Documentation and tutorials are available at: https://sgs.bioinfotoolkits.net/home.

## Author Contributions

T. X. and Y.W. conceptualized the project. Y.W. supervised the project. T.X., H.S., F.L. and J.L. designed and wrote the software code. T.X. and H.S. conducted software testing and analysis. T.X. and H.S. performed data collection and labeling. Y.W. finalized the manuscript with input from all authors. All authors read and approved the final manuscript.

## Acknowledgements

We thank Ling Xu at China Agricultural University, Zhaoyuan Wei and Guoqing Zhang at Southwest University for critical reading of the manuscript. National Natural Science Foundation of China [U21A20248]; National Natural Science Foundation of China [32000340]; Fundamental Research Funds for the Central Universities [XDJK2019TJ003]. Funding for open access charge: National Natural Science Foundation of China.

