## Supplementary information for "Empowering Integrative and Collaborative Exploration of Single-Cell and Spatial Multimodal Data with SGS"

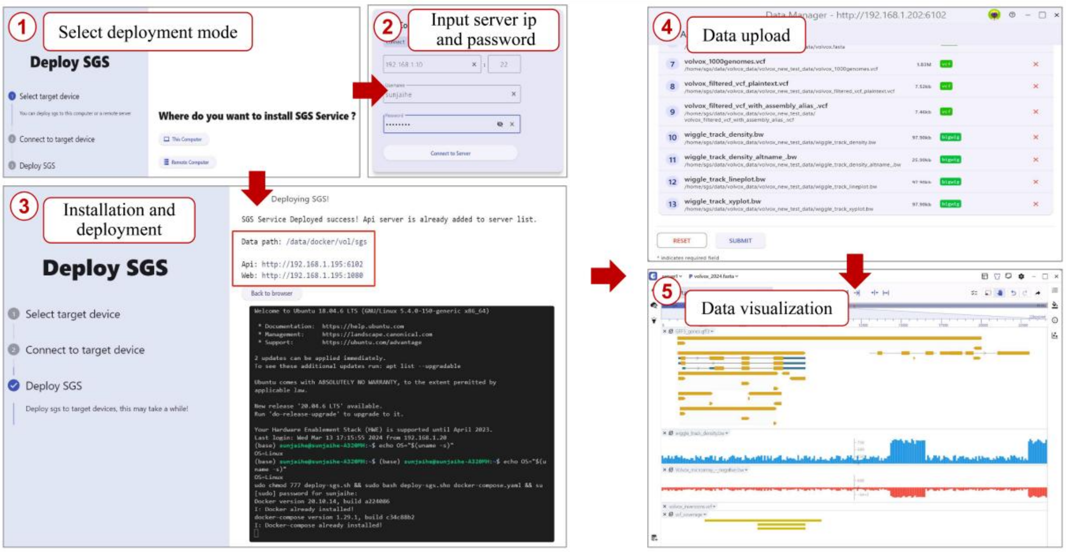


**Supplementary Figure 1:** Rapid installation and deployment of SGS on local or remote servers. After downloading the SGS client, users can deploy the SGS visualization server using the user-friendly GUI interface. ① In this interface, you'll be presented with two installation modes to choose from: local installation (recommended for deployment on a local computer) or remote installation (ideal for deployment on a remote server). ② Select the desired mode and provide essential information such as host IP address, username, password, and port. ③ During the deployment process, a real-time progress interface will keep you informed about the status of the deployment. Once deployment is successful, click 'Back to Browser' to manage data. ④ Users can add multiple files by selecting a folder or individual files for submission. After adding the data, you can access the data list interface to view the status of the added data, perform queries, and delete specific files. ⑤ Finally, users can explore the added data in the browser interface.

| | Cellxgene | iSEE | Loom-Viewer | scSVA | SCope | Single Cell Explorer | UCSC Cell Browser | Loupe Browser | ST Viewer | TissUUmaps3 | AtlasXplore | Vitessce | SGS |
|---|---|---|---|---|---|---|---|---|---|---|---|---|---|
| **cell selection and export** | ✓ | ✓ | | ✓ | ✓ | ✓ | ✓ | ✓ | ✓ | ✓ | ✓ | ✓ | ✓ |
| **Zoom in/out** | ✓ | ✓ | | ✓ | ✓ | | ✓ | ✓ | | ✓ | | ✓ | ✓ |
| **Multiple embeddings** | ✓ | ✓ | ✓ | | ✓ | | ✓ | ✓ | ✓ | ✓ | ✓ | ✓ | ✓ |
| **Highlight gene expression** | ✓ | ✓ | | ✓ | ✓ | ✓ | ✓ | | ✓ | ✓ | ✓ | ✓ | ✓ |
| **Highlight metadata** | ✓ | ✓ | ✓ | ✓ | | ✓ | ✓ | | ✓ | ✓ | ✓ | ✓ | ✓ |
| **spatial expression visualization** | ✓ | | | | | | ✓ | ✓ | ✓ | ✓ | ✓ | ✓ | ✓ |
| **tissue image visualization** | ✓ | | | | | | ✓ | ✓ | ✓ | ✓ | | ✓ | ✓ |
| **Flexible layout** | | | | | | | | | | | | ✓ | ✓ |
| **Split screen support** | | | | | | | ✓ | ✓ | | | ✓ | ✓ | ✓ |
| **multi-feature comparison** | | | | | | | | | | | ✓ | | ✓ |
| **multi-slice visulization** | | | | | | | | | | ✓ | | | ✓ |
| **Synchronization between windows** | | | | | | | | | | | ✓ | ✓ | ✓ |
| **genome-mapping track** | | | | | | | | ✓ | | | ✓ | ✓ | ✓ |
| **Web page loads fast** | ✓ | | ✓ | ✓ | ✓ | ✓ | ✓ | | | ✓ | ✓ | ✓ | ✓ |
| **Platform** | Python | R | Python | R | Python | Python | Python | Desktop | C++ | Javascript | Javascript | Javascript | Python |
| **Web Sharing** | ✓ | ✓ | ✓ | ✓ | ✓ | ✓ | ✓ | | | ✓ | ✓ | ✓ | ✓ |
| **Interactivity** | ✓ | ✓ | | | ✓ | ✓ | ✓ | | ✓ | ✓ | ✓ | ✓ | ✓ |
| **Cloud support** | ✓ | | | ✓ | ✓ | | ✓ | | | ✓ | ✓ | ✓ | ✓ |
| **Session URL sharing** | ✓ | | | | | | ✓ | | | | | | ✓ |
| **loupe** | | | | | | | | ✓ | | | | | |
| **anndata/anndata.zarr** | ✓ | | | ✓ | | ✓ | ✓ | | | | | ✓ | ✓ |
| **mudata/mudata.zarr** | | | | | | | | | | | | ✓ | ✓ |
| **Loom** | | | ✓ | ✓ | ✓ | ✓ | ✓ | | | | | | |
| **SCE** | | ✓ | | | | | | | | | | | |
| **Seurat** | | | | | | ✓ | ✓ | | | | | | |
| **csv/txt** | | | | ✓ | | ✓ | ✓ | | ✓ | ✓ | ✓ | ✓ | |
| **GFF** | | | | | | | | | | | | | ✓ |
| **VCF** | | | | | | | | | | | | | ✓ |
| **BED** | | | | | | | | | | | | ✓ | ✓ |
| **BigWig** | | | | | | | | | | | ✓ | ✓ | ✓ |
| **MelthyC** | | | | | | | | | | | | | ✓ |
| **Longrange** | | | | | | | | | | | | | ✓ |
| **BigInteract** | | | | | | | | | | | | | ✓ |
| **HIC** | | | | | | | | | | | | | ✓ |
| **GWAS** | | | | | | | | | | | | | ✓ |

**Supplementary Table 1:** Comparison of SGS with existing mainstream single-cell and spatial multi-omics visualization tools: Cellxgene, iSEE, Loom-Viewer, scSVA, SCope, Single Cell Explorer, UCSC Cell Browser, Loupe Browser, ST Viewer, TissUUmaps3, AtlasXplore, Vitessce. The orange section primarily highlights the fundamental visualization features such as cell selection, gene expression, and spatial slice display. The light blue section focuses on advanced visualization features including multiple features comparison, multi spatial slices comparison, multi-panel coordination, and genome-mapped data visualization. The light green section compares these tools in terms of loading speed, programming language, visualization sharing, and local deployment. The light yellow section provides an in-depth comparison of the supported file formats across these tools.

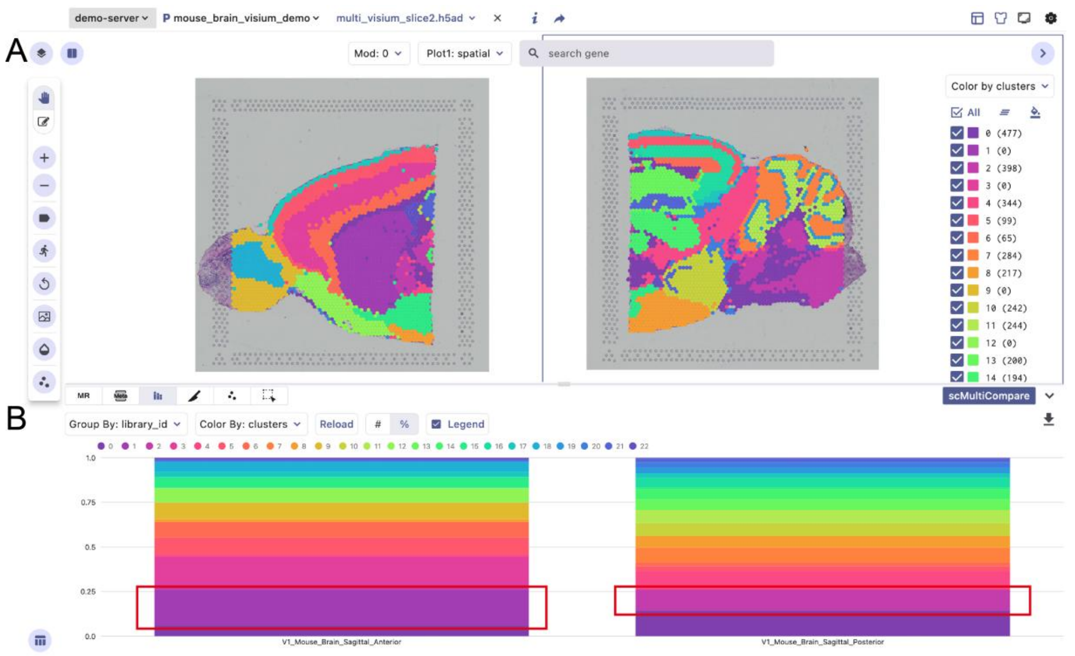


**Supplementary Figure 2:** The SC module displays 10x Visium data of the anterior and posterior mouse brain. (**A**) The left and right panels presents the cell cluster of the anterior and posterior brain spatial slices respectively. (**B**) The cell proportion plots were generated for mouse brain, grouped by sample ID.

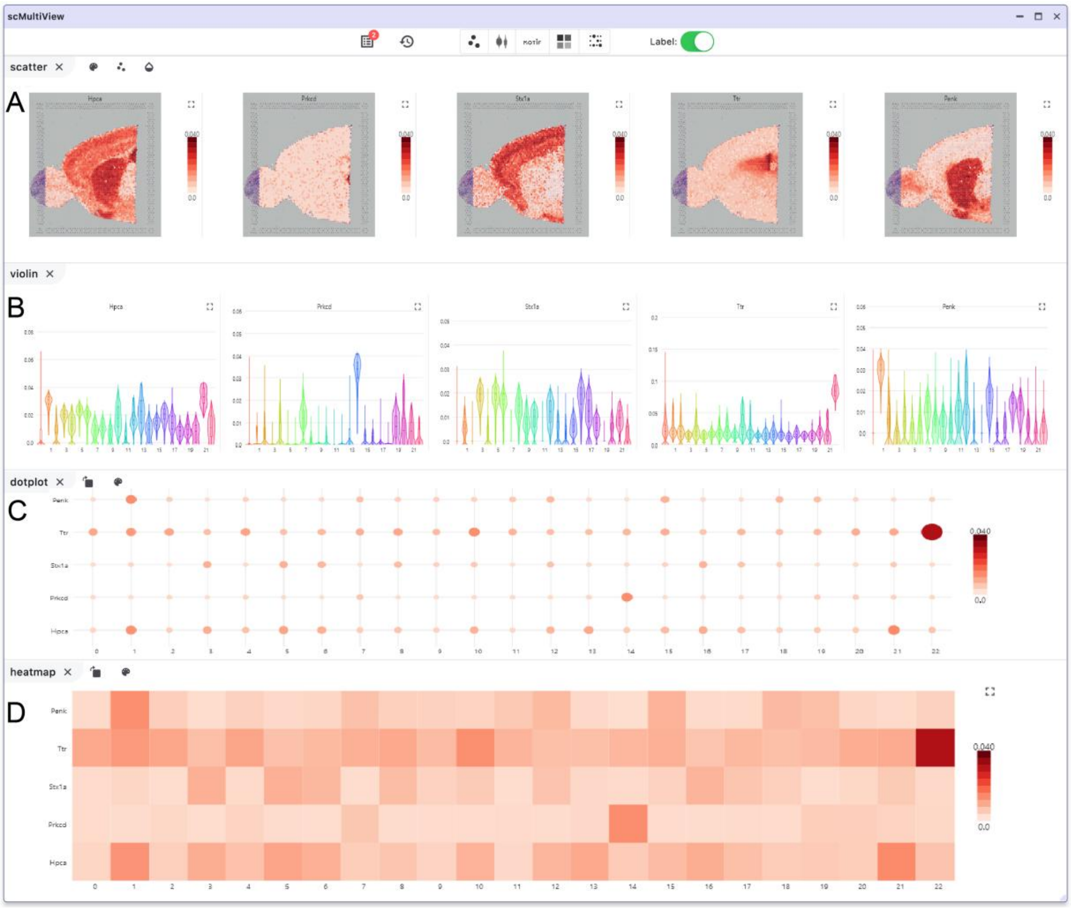


**Supplementary Figure 3**: Visualizing development-related genes (*Hpca*, *Prkcd, Stx1a*, *Ttr*, *Penk*) in 10x Visium mouse brain data using the scMultiview component. The comparative analysis is based on cell clusters and showcases spatial feature plot (**A**), Violin plot (**B**), dot plot (**C**), and Heatmap (**D**) for the selected genes.

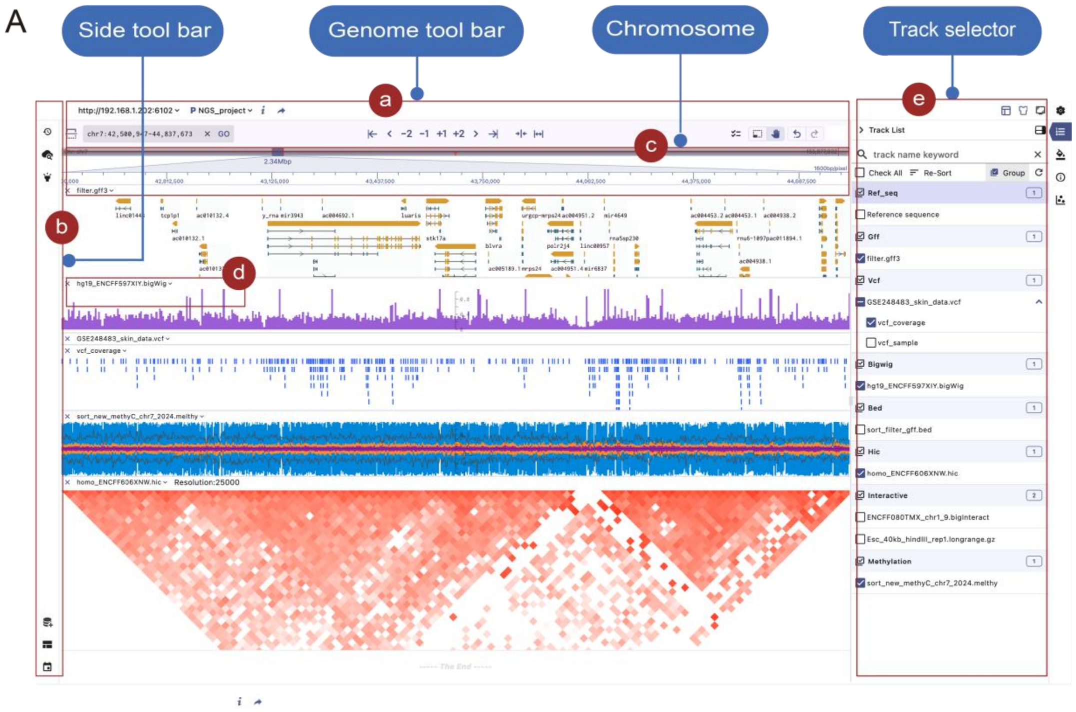


**Supplementary Figure 4**: The novel genome browser framework of SGS. **(A)** The linear genome visualization panel consists of three main components: the genome toolbar (**a**), side toolbar (**b**) and main visualization panel (**c**). Additionally, track setting (**d**) and track selector (**e**) are also available for track selection and customization, respectively.

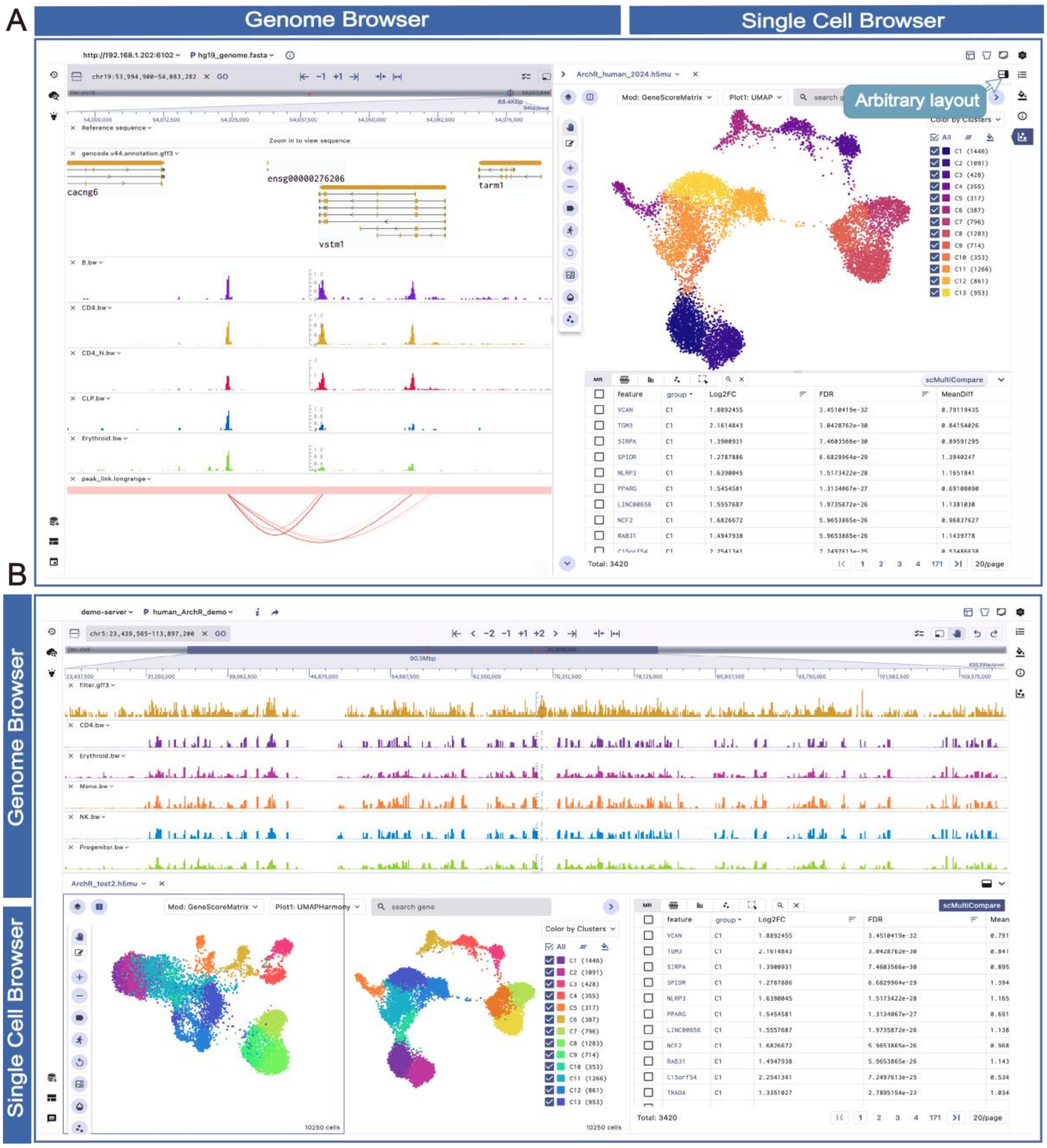


**Supplementary Figure 5:** Flexible interface layout in SG visualization module. The depicted figure showcases the SG_H and SG_V layout modes of the SG module. (**A**) The SG_H layout mode emphasizes assisting users in comparing multiple tracks in specific genomic regions, such as chromosome gene annotations and epigenetic modification signals, with single-cell omics information. (**B**) On the other hand, the SG_V layout mode vertically arranges the single-cell data and genome information, allowing users to obtain a view with fewer genome track details across the entire chromosome.

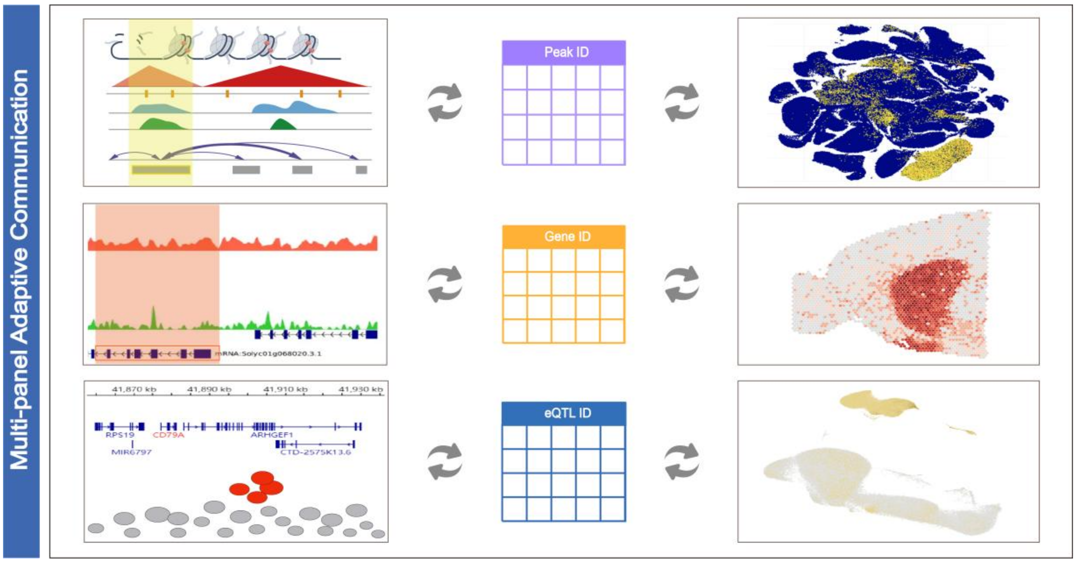


**Supplementary Figure 6**: The multiple-panel adaptive communication mechanism, showing interaction between the genome framework and single-cell components. Clicking on features like peak ID, gene ID, or sc-eQTL ID in the marker table highlights the corresponding genomic region and its single-cell expression distribution in both panels.

### User Case

#### Single-cell eQTL Data Visualization

The SGS Multi-Omics Browser provides researchers with a powerful tool to visually explore sc-eQTL (single-cell expression quantitative trait loci) data, providing valuable insights into the impact of gene regulation on cellular function and disease mechanisms. The illustration showcases sc-eQTL data from the OneK1K research, including 982 individuals[1]. The genome browser presents multiple Gaws tracks of sc-eQTLs across various cell types. Users can easily locate specific sc-eQTL regions by entering gene symbols (e.g. *BLK*) or genomic positions (e.g. chr8) (**Supplementary Figure 7 (a,b)**). The single-cell visualization component displays UMAP of 14 major cell types (**Supplementary Figure 7 (c)**), while the interactive table at the bottom lists marker genes and sc-eGenes (genes significantly associated with sc-eQTLs) related to specific cell types (**Supplementary Figure 7 (d)**). Clicking on a gene in the marker table allows users to examine its expression patterns across cell types and navigate to the associated sc-eQTL region in the genome browser.

For example by clicking on the "*BLK*" gene in the marker table, users can explore cell expression heterogeneity and navigate to the corresponding genomic region in the genome browser. In the single-cell panel, we can observe it specific high expression in Bmen cells. In the genome Browser panel, *BLK*-specific sc-eQTLs are highlighted in red, while other sc-eQTLs are marked in gray. Users can access details such as *BLK* sc-eQTL site ID, p-value, and more. Here, we observe that many eSNPs associated with the "*BLK*" gene (linked to autoimmune diseases) show correlations with BLK expression in CD4 NC, CD8 ET, CD8 NC, B Mem, and B IN cells. Notably, among the associated eQTL loci, the rs2736336 locus was identified. Previous studies have reported that the *rs2736336* variant leads to differential expression of *BLK* in B Mem cells, indicating its potential role in inter-individual variability of B lymphocyte tolerance.

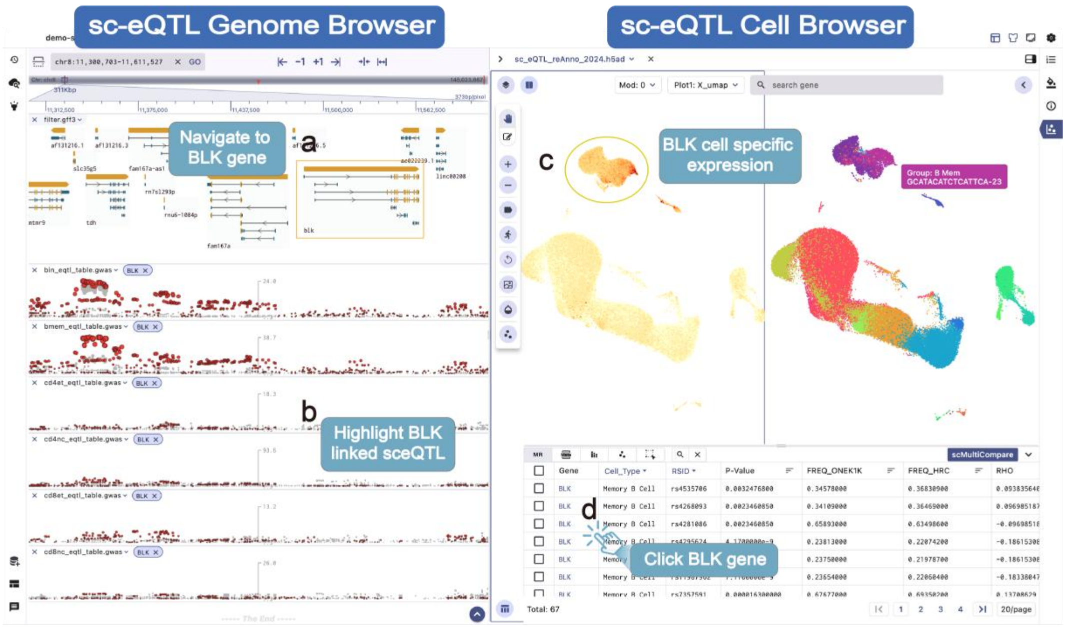


**Supplementary Figure 7:** The screenshot of OneK1K sc-eQTL visualization demo. The panel consists of two components: the genome framework and the single-cell component. The genome framework provides insights into the structure of the *BLK* gene region (**a**) and highlights cell type-specific sc-eQTL loci. The significance of each loci is represented by the size and height of the points, with sc-eQTLs associated with the "*BLK*" gene highlighted by black circles (**b**). The single-cell component displays the UMAP of 14 cell clusters and the expression pattern of the *BLK* gene across different cell populations (**c**). The marker table displays essential information such as the p-value, q-value, log fold change (logFC), and other relevant details of the marker gene (**d**).

### ME11 Spatial-ATAC-seq Visualization

By exploring the spatial chromatin accessibility landscape during mouse embryo development, the SGS browser showcases the Spatial-ATAC-seq data integrative visualization. The genome browser supports selecting and grouping chromatin accessibility signal tracks[2]. By clicking the navigation button of a target gene, users can swiftly locate the genomic region and visualize the distribution trends of accessibility signals among different cell clusters (**Supplementary Figure 8 (a)**). Additionally, SGS allows selecting target genes from the marker table to visualize spatial expression and tissue slice. This integrated visualization approach enables users to quickly observe the epigenomic signals and spatial expression differences of target genes at the single-cell level, linking epigenetic signals with spatial slice context information for a more comprehensive understanding of gene regulation mechanisms.

Coupled with tissue slice information, we identified four clusters with distinct spatial patterns in the spatial ATAC sequencing data of 11-day-old mouse embryos (**Supplementary Figure 8 (e)**). Among these clusters, the "*RARG*" gene (associated with prominent facial features and limb development in the embryo) is extensively activated in Cluster 4 (**Supplementary Figure 8 (c)**), indicating a potential crucial role of the cluster in limb bud development, skeletal growth, and matrix balance. By clicking the rapid navigation button for the *RARG* gene (**Supplementary

**Figure 8 (d)**), users can navigate to the genomic region and visualize the *RARG* gene exhibits higher chromatin accessibility signals in Cluster 4 compared to other clusters (**Supplementary Figure 8 (a,b)**). SGS providing a powerful tool for better understanding the gene regulation mechanisms and expression patterns during mouse embryo development.

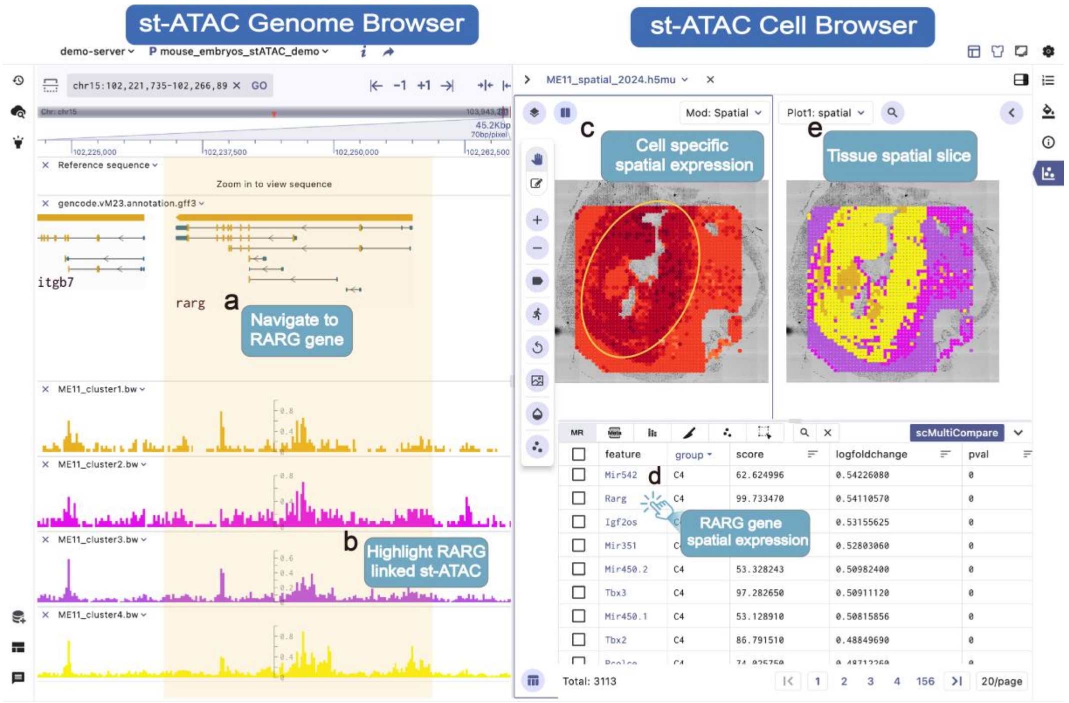


**Supplementary Figure 8:** Integrative visualization of spatial ATAC dataset in 11-day mouse embryos. The genome view (**a,b**) shows the structure of the *RARG* gene and chromatin accessibility signal across different cell clusters, highlighting cluster 4 as having the highest signal. The spatial view (**c-e**) reveals the cell types and spatial expression differences of the *RARG* gene.

A

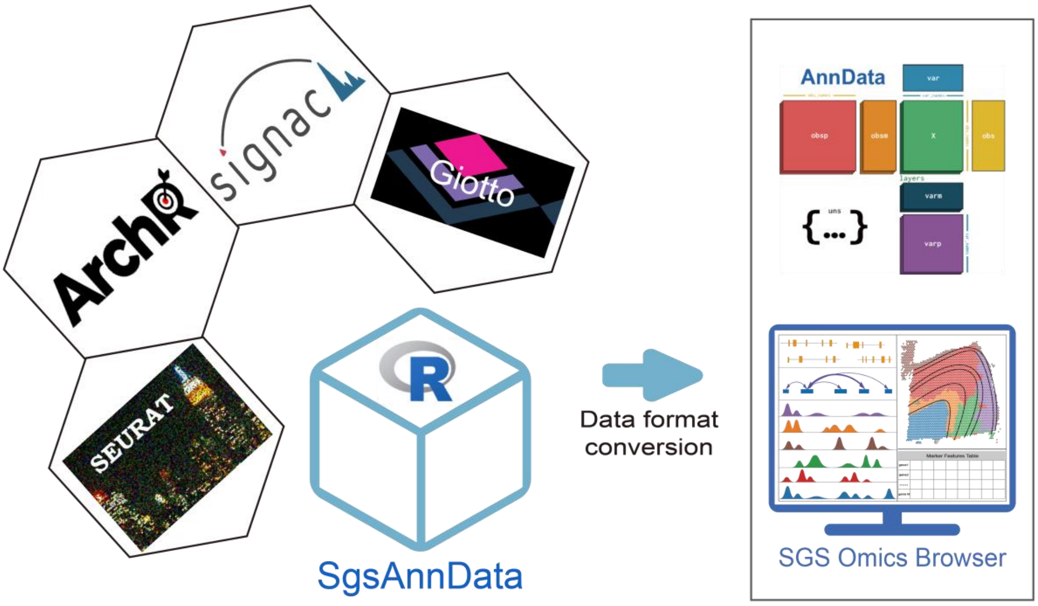


B

#### Seurat to AnnData

##### Single-Cell Transcriptome Object Conversion

When Seurat contains multiple assays, users can provide multi assay names they wish to export. This will automatically generates the h5ad for each assay type, like RNA.h5ad,integration.h5ad etc. We also offer the flexibility for users to output marker dataframes as a list, with the corresponding information stored in **adata.uns['markers']**.

```
library(Seurat)
library(anndata)
SeuratToAnndata(object=scRNA,
                outpath="/test_adata",
                assays=c("RNA", "SCT"),
                groups = NULL,
                reductions = c("tsne","umap"),
                markersDF = list("RNA"=rna_marker_df, "SCT"=sct_marker_df))
```

#### Signac to Anndata

We recommend exporting **peak matrix**, **genescore matrix**, and **motif matrix** from single-cell ATAC data analysis as h5ad files. If desired, these files can be further converted into Mudata, which offers a comprehensive and annotated multimodal dataset structure. Additionally, users can use **export_pwm** to output results from co-accessibility analysis and motif enrichment analysis, respectively. Users can access motif-related information via **adata.uns.["motifs"]**.

```
SignacToAnndata(object=scATAC,
                outpath = "/test_adata",
                assays=c("RNA", "Peaks", "Motif"),
                assay.types=("gene","peak","motif"),
                markersDF = list("RNA"=gene_marker_df, "Peaks"=marker_peak_df, "Motif"=marker_motif_df)
                groups = c("clusters","sample", "age"),
                reductions = c("umap","tsne"),
                export_pwm = TRUE)   # to export motif pwm
```

**Supplementary Figure 9:** (**A**) SgsAnnData is an R package simplifies the conversion of single-cell analysis object from popular tools such as Seurat, Giotto, Signac, and ArchR into the AnnData format. This format can be directly visualized in SGS. (**B**) The SgsAnnData package is used to convert Signac objects into multiple h5ad files, which are then stored in MuData format. To perform this conversion, users need to utilize the SignacToAnndata function and provide parameters such as assay, metadata, reduction, motifs, markers etc. Individual h5ad objects are created for each assay. Subsequently, the mudata package is employed to create a MuData file, which can be directly imported into the SGS multi-omics browser for visualization.

**Supplementary Table 2:** Summary of datasets used in this manuscript

| Figure | Data URL | PMD |
|---|---|---|
| Figure 4A | https://pubmed.ncbi.nlm.nih.gov/31792411/ | PMC7258684 |
| Figure 3A | https://db.cngb.org/stomics/mosta/ | PMID: 35512705 |
| Figure 3A | https://genome.ucsc.edu/s/brianpenghe/scATAC_fetal_lung20211206 | PMID: 36493756 |
| Figure 5A | https://cells.ucsc.edu/?ds=brain-epigenome+human-brain-m3c | / |
| Figure 5 B | https://www.10xgenomics.com/datasets?menu%5Bproducts.name%5D=Spatial%20Gene%20Expression | / |
| Supplementary Figure 3 | https://epigenomegateway.wustl.edu/browser/ | / |
| Supplementary Figure 5 | https://cellxgene.cziscience.com/collections/dde06e0f-ab3b-46be-96a2-a8082383c4a1 | PMID: 35389779 |
| Supplementary Figure 6 | https://www.ncbi.nlm.nih.gov/pmc/articles/PMC9452302/ | PMCID: PMC9452302 |
